# Factor Analysis of Multimodal MRI, Biofluid and Vascular Health Biomarkers Reveals Latent Constructs of Brain Health

**DOI:** 10.1101/2025.05.06.652544

**Authors:** Ella Rowsthorn, Ming Ann Sim, William T. O’Brien, Stuart J. McDonald, Katherine Franks, Benjamin Sinclair, Trevor T.-J. Chong, Stephanie Yiallourou, Marina Cavuoto, Lucy Vivash, Terence J. O’Brien, Xingfeng Shao, Danny J.J. Wang, Meng Law, Ian H. Harding, Matthew P. Pase

**Author notes:** Correspondence: Assoc/Prof. Matthew P. Pase, School of Psychological Sciences, Monash University, 18 Innovation Walk, Clayton VIC 3168, Australia, E. These authors contributed equally (Ian H. Harding, Matthew P. Pase).

## Abstract

Individual imaging and fluid biomarkers provide insights into specific components of brain health, but integrated multimodal approaches are necessary to capture the complex, interrelated biological systems that contribute to brain homeostasis and neurodegenerative disease.

Using data from the Brain and Cognitive Health (BACH) cohort study (N=127; mean age=67 years, 68% women), we performed an exploratory factor analysis to identify latent constructs of brain health. We included multimodal neurovascular imaging markers, brain atrophy metrics, plasma Alzheimer’s disease (AD) biomarkers and cardiovascular risk factors. Five constructs emerged: “Brain & Vascular Health” (greater hippocampal volume, basal ganglia enlarged perivascular spaces [ePVS], cerebral blood flow and HDL cholesterol; lower ventricle volume and BMI); “Structural Integrity” (greater cortical thickness, fractional anisotropy and basal ganglia ePVS); “Fluid Transport” (greater white matter ePVS and Free Water); “AD Biomarkers” (higher phosphorylated tau [pTau]181 and pTau217; lower amyloid-beta 42/40 ratio); and “Neuronal Injury” (higher glial fibrillary acidic protein and neurofilament light chain). All constructs were associated with age (β=-0.70–0.39, *p*≤.014), except for Fluid Transport (*p*>.05). Brain & Vascular Health and Structural Integrity (*partial r*=.305, *p*<.001), and AD Biomarkers and Neuronal Injury (*partial r*=.248, *p*=.005) were positively correlated. Only Brain & Vascular Health was associated with global cognition (β=0.27, SE=0.13, *p*=.043).

These findings provide a data-driven framework for examining distinct constructs underlying vascular health, fluid regulation and neurodegenerative pathology. We demonstrate the utility of using multiple biomarkers to probe these biological systems, paving the way for future research to explore how these systems change across diverse neurodegenerative conditions.

## Introduction

The brain relies on several interconnected systems to maintain homeostasis and support cognitive function, including the neurovascular unit for nutrient delivery and the fluid transport system for removing metabolic waste^1,2^. Growing evidence indicates that neurovascular and fluid transport systems become dysfunctional in the earliest stages of many neurodegenerative diseases, including Alzheimer’s disease (AD), with these changes often preceding the appearance of clinical symptoms and significant brain atrophy^3–5^. It is evident that the dynamic interplay between these systems is essential for maintaining brain resilience against aging and pathology^6^ but these nuanced inter-relationships present significant challenges for assessing the overall health of the brain directly.

As the repertoire of biomarkers frequently captured in aging and neurodegenerative research has expanded, there is emerging recognition that a combination of markers may provide a more nuanced and comprehensive assessment of brain health and neurodegenerative mechanisms. For instance, interdependencies between different MRI measures of neurovascular health provide broader insights into neurobiological processes that become impaired in neurodegenerative disease, beyond inferences that can be drawn from individual metrics alone^7,8^. Similarly, the ratio or combination of multiple plasma or cerebrospinal fluid biomarkers has been demonstrated to better stage AD severity^9^, and is useful in differentiating AD from other diseases^10^. However, the synergistic potential of combining information across multiple data modalities (e.g., imaging, biofluid, lifestyle) for detection of more subtle changes in brain health, such as in normative aging or early phases of pathology, remains largely unexplored.

In the case of AD, pathology risk and severity are influenced by a diverse range of factors, including lifestyle, cardiovascular health, and efficiency of the brain’s neurovascular fluid transport and waste clearance system^4,11,12^. A recent review has emphasised that these distinct factors are also highly inter-dependent, where dysfunction of one system may influence dysfunction in another and interact with neurodegenerative pathology^13^. For example, impaired fluid transport through the brain’s waste clearance system may promote the accumulation of fibrillar amyloid-beta (Aβ) and phosphorylated-tau (pTau) which are hallmark pathological markers of AD^14,15^. Simultaneously, modifiable cardiovascular risk factors, such as elevated LDL cholesterol, hypertension and obesity significantly contribute to late-life dementia risk^16,17^, and are linked the neurovascular dysfunction seen in AD, such as reduced cerebral blood flow, impaired neurovascular coupling and perivascular enlargement^18–21^.

The complex interactions that exist within and between the cardiovascular, neurovascular, and brain fluid regulation systems in both aging and neurodegenerative disease drive the need for a more holistic research approach. Individual markers of brain health provide useful insight into specific elements or sub-components of these systems. However, identifying latent constructs by capturing the shared variance across multiple neurovascular, cardiovascular and neurodegenerative markers may better probe the core and potentially distinct biological constructs underlying aging and pathological processes and offer a more comprehensive and sensitive measure of brain health.

In this study, we conducted an exploratory factor analysis to identify the latent constructs underlying multiple neurovascular imaging markers, brain atrophy metrics, plasma biomarkers and cardiovascular risk factors in mid-late life adults from the community. We aimed to derive a set of weighted composite measures from the identified factors and investigated how these composites that reflect different aspects of neural health are associated with age and cognitive function. By taking a more comprehensive approach, our work aimed to facilitate the development of more sensitive and integrated composite measures for studying the aging brain, with the hypothesis that these composites would meaningfully capture distinct aspects of neurovascular health or neurodegenerative mechanisms.

## Method

### Study Participants

The Brain and Cognitive Health (BACH) cohort is a prospective study that recruited older adults from the community. Participants underwent blood sample collection, over-the-phone cognitive assessments, an in-person neuropsychological battery, and an MRI scan. To be eligible, participants were 55-80 years of age and were without self-reported dementia, significant neurological disease (e.g. Parkinson’s disease, epilepsy) or history of disabling stroke. Exclusion criteria included contraindications to MRI, and evidence of moderate-severe cognitive impairment (assessed with the TELE adapted for Australians^22^). The BACH cohort study was approved by the Alfred Health Ethics Committee (project 78642) and each participant provided written informed consent. This study used cross-sectional data from the first 149 participants, obtained from 2022 to 2023.

### Plasma Biomarkers of AD

All participants underwent a fasted blood sample collection for both immediate clinical blood-panel analysis and storage for subsequent biomarker analysis. Blood for plasma biomarker analysis was collected in EDTA tubes with prostaglandin E1, then aliquoted and stored at -80 degrees.

Plasma concentrations of fluid biomarkers were quantified using the SIMOA HD-X (Quanterix, Boston, USA) in the Department of Neuroscience, School of Translational Medicine, Monash University, in accordance with manufacturer’s instructions. Glial fibrillary acidic protein (GFAP), neurofilament light chain (NfL), Aβ40 and Aβ42 were quantified using the SIMOA Human Neurology 4-Plex E assay; pTau-181 was quantified using the SIMOA pTau-181 Advantage V2.1 assay; and pTau-217 was quantified using the ALZpath pTau-217 Advantage PLUS assay. A single sample was analysed on all plates as a control. All samples had concentrations above the above the manufacturer’s analytical lower limit of quantification (LLOQ) for each biomarker, which was 2.89 pg/mL for GFAP, 0.40 pg/mL for NfL, 1.02 pg/mL for Aβ40, 0.378 pg/mL for Aβ42, 2.00pg/mL for pTau-181 and 0.00326pg/mL for pTau-217. The mean coefficient of variation for all biomarkers were below 10%: 4.6% for GFAP; 4.3% for NfL; 2.6% for Aβ40; 2.7% for Aβ42; 7.8% for pTau-181; 5.8% for pTau-217.

The ratio of Aβ42 to Aβ40 (Aβ42/40) has been shown to be particularly sensitive to pathological changes of amyloid levels in the earliest stages of AD, before overt cognitive impairment^23–25^, and more robust to individual variations in total Aβ^26^. Given this, we used the Aβ42/40 ratio in the present analysis rather than each measurement separately.

### MRI Markers of Neurovascular and Brain Health

All participants were scanned on a 3T Siemens Prisma MRI machine using a 64-channel head coil. The scanning protocol has been described previously^7^, which included T1-weighted, T2-weighted fluid-attenuated inversion recovery (FLAIR), multi-shell diffusion-weighted imaging, pseudo-continuous arterial spin labelling (pCASL) and diffusion-prepared pCASL (DP-pCASL).

Measures of neurovascular integrity and fluid transport, including whole-brain blood-brain barrier water exchange rate (BBB k_w_), gray matter cerebral blood flow (CBF), white matter isotropic diffusion volume fraction (‘Free Water’), white matter hyperintensity (WMH) volume and MRI-visible enlarged perivascular space (ePVS) volume were processed as reported previously (Figure 1)^7^. Briefly, estimated total intracranial volume, gray matter and white matter segmentations were derived from Fastsurfer^27^ (v1.0.0, e4ed6f7); BBB kw was derived from DP-pCASL using the LOFT toolbox^28^; CBF was derived from pCASL using FSL BASIL^29^; Free Water was derived from diffusion-weighted imaging, pre-processed through QSIPrep^30^ (version 0.14.3, based on Nipype 1.6.1) and analysed with three-compartmental NODDI modelling^31^; WMH was segmented on FLAIR images automatically using an in-house nnUnet model; and ePVS was segmented on T1-weighted images using PINGU^32^. For the present study, given that ePVS regions are differentially associated with vascular and amyloid angiopathy related diseases^33,34^, ePVS segmentations were further sectioned into ‘normal appearing white matter’ (WM) and ‘basal ganglia’ (BG) (i.e., the caudate, putamen, pallidum, accumbens, thalamus, ventral diencephalon) regions according to the segmentations from Fastsurfer. These ePVS volumes were divided by the total volume of the normal appearing white matter and BG gray matter, respectively, deriving WM and BG ePVS volume fraction.

**Figure 1:**
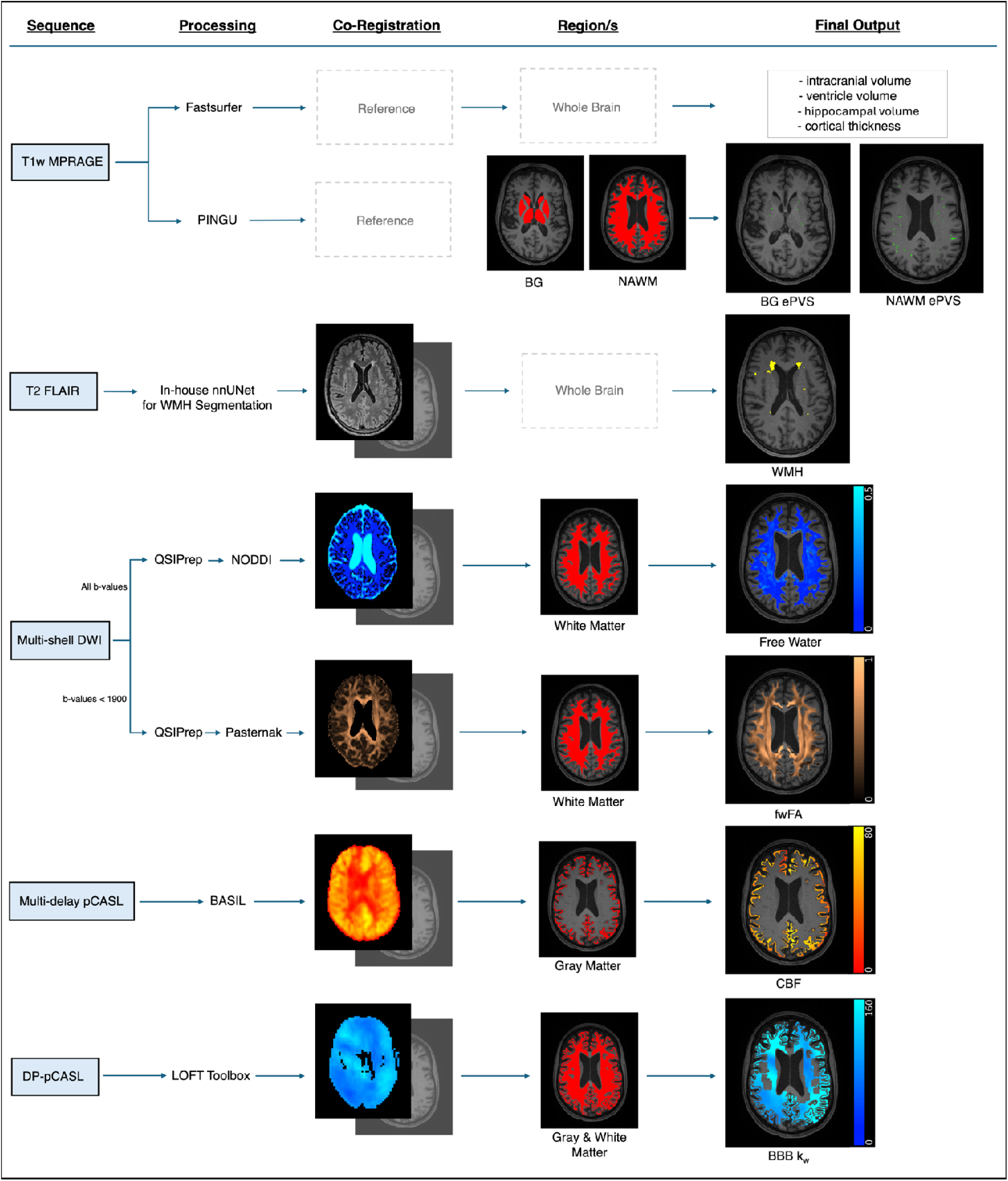
MRI Processing Flow Diagram. All participants underwent a Siemens 3T MRI scan. T1w images were processed through Fastsurfer to segment regions of interest and quantify **brain volumes**. Separately, enlarged perivascular spaces (**ePVS**) were segmented via PINGU from the T1w images and volume fractions were quantified within basal ganglia (BG) and normal-appearing white matter (NAWM, i.e. excluding white matter hyperintensities (WMH)). **WMH** were segmented on FLAIR images by an in-house nnUNet trained on study data, reviewed for accuracy. All DWI images were pre-processed through QSIPrep and analysed with NODDI three-compartmental modelling to derive **Free Water**. An upper threshold of 0.5 was applied to Free Water maps to minimise erroneous partial-volume effects. Since Pasternak bi-tensor modelling is affected by non-Gaussian effects in b-values greater than 1900s/mm , DWI images with b-values 0-1500s/mm were separately processed through QSIPrep and analysed with Pasternak to derive Free Water-corrected fractional anisotropy (**fwFA**). Pseudo-continuous arterial spin labelling (pCASL) data was pre-processing and analysed with FSL’s Bayesian Inference for Arterial Spin Labelling MRI (BASIL) to derive cerebral blood flow (**CBF**). Lastly diffusion-prepared (DP-pCASL) was processed through the Laboratory of Functional MRI Technology (LOFT) toolbox for image reconstruction, pre-processing and to derive blood-brain barrier water exchange rate (**BBB k_w_**).

In addition to the measures of neurovascular integrity and fluid transport, we also calculated measures of macro- and micro-structural integrity. Ventricle volume, hippocampal volume and cortical thickness were also derived from Fastsurfer analysis, with the volume measures divided by total estimated intracranial volume. We also calculated Free Water-corrected fractional anisotropy (fwFA), a sensitive measure of white matter microstructural integrity that minimises noise caused by isotropic diffusion. To quantify fwFA we firstly removed image volumes acquired with a b-value greater than 1900s/mm^2^ from the diffusion data to minimise non-Gaussian effects, as recommended for bi-tensor models of Free Water quantification^35^. We then pre-processed the diffusion data through QSIPrep and quantified fwFA using the model described by Pasternak et al^36,37^. The resulting images were linearly registered to the participant’s T1-weighted image using Advanced Normalization Tools^38^ (ANTs), where the whole brain fwFA intensity map was then linearly transformed to the T1-weighted space using this registration. Possible fwFA values range from 0 to 1, where higher values represent greater white matter microstructural integrity (i.e., more anisotropic diffusion).

### Cardiovascular Risk Factors

Participants underwent in-person assessment of their height, weight and blood pressure. Body-mass index (BMI) was calculated as weight divided by height squared. Average systolic blood pressure was calculated as the average of three supine assessments taken after 5-minutes of rest. LDL and HDL cholesterol levels were obtained from a freshly drawn blood sample, taken in the morning following an overnight fast.

### Cognitive Assessment

Participants were administered over-the-phone and in-person cognitive assessments that evaluated performance across several cognitive domains, including: Prose Passages delayed recall^39^ (phone version) to assess short-term memory; the Wechsler Adult Intelligence Scale (WAIS-IV) Similarities^40^ (phone version) to assess reasoning; Sydney Language Battery (SYDBAT) naming task^41^ to assess language; Controlled Oral Word Association Test (COWAT) Verbal Fluency^42,43^ to assess executive function; Trail Making Test A and B^44^ to assess processing speed and executive function; the Wechsler Memory Scale (WMS-IV) Visual Reproduction^45^ to assess spatial memory; the Hooper Visual Organisation Test^46^ for visuo-spatial processing; and The Awareness of Social Inference Test (TASIT)^47,48^ to assess social cognition. For most participants, over-the-phone and in-person cognitive assessments were completed within two weeks of each other (mean difference was 8 days) except for a handful of subjects due to re-scheduling (n=11, only one participant exceeded four weeks).

A principal components analysis of all cognitive tests was used to define a global composite score. Trail Making Task A and the TASIT did not load onto the main factor and were removed from the model. Cronbach’s alpha testing verified that the other seven tests should remain to maximise factor reliability: Prose Passages delayed, Similarities, SYDBAT Naming, Visual Reproductions delayed recall, Trail Making Test B, Verbal Fluency, the Hooper Visual Organisation Test (analysis detailed in the Supplement). A global cognition composite was created by summing the z-scores of these seven variables (Trail Making Test B was reverse coded) and dividing by seven. Higher scores indicate better global cognitive performance. Due to the presence of a single extreme outlier, the global cognition composite scores were Winsorized to the 1^st^ and 99^th^ percentiles to mitigate outlier influence while preserving the overall distribution of the data.

Additionally, the Mini-Mental State Examination (MMSE)^49^ and Clinical Dementia Rating scale (CDR)^50,51^ were administered to assess global orientation and dementia status respectively, used to describe the sample.

#### Data Analysis

To identify potential latent constructs within the data, an exploratory factor analysis was conducted. Unlike principal components analysis, where the primary purpose is to reduce the dimensionality of the data by extracting components that maximally explain the total variance observed (including unique variance), exploratory factor analysis seeks to identify underlying latent factors by isolating the *shared* variance among observed variables – an inherently agnostic and data-driven approach.

Firstly, to determine the number of sufficient factors, a parallel analysis was conducted on the standardised variables, including the ten imaging, six plasma biomarkers and four cardiovascular risk variables (Table 1). Models were considered reliable if the Tucker-Lewis Index of Factoring Reliability was greater than 0.90 and demonstrated goodness of fit when the RMSEA index was below 0.05. Variables with factor loadings greater than 0.30 were considered meaningful indicators of the latent construct and were used to create scaled, weighted composite scores for each construct (i.e. construct composites).

**Table 1.** Cohort Summary.

| **Sample Characteristics**<br>(N=127) |  |
| --- | --- |
| **Demographics** |  |
| Women (n, %) | 86 (67.7%) |
| Age, years | 66.82 ± 5.29 |
| Education, years | 17.32 ± 3.28 |
| **Cardiovascular Health** |  |
| Systolic Blood Pressure, mmHg | 128.51 ± 16.28 |
| HDL Cholesterol, mmol/L | 1.67 ± 0.39 |
| LDL Cholesterol, mmol/L | 3.21 ± 0.91 |
| Body Mass Index | 26.23 ± 4.77 |
| **Cognition** |  |
| MMSE, score out of 30 | 28.57 ± 1.44 |
| CDR Sum of Boxes ≥ 0.5 (n, %) | 22 (17.3%) |
| Prose Passages Delayed, % correct | 0.40 ± 0.14 |
| Similarities, score out of 36 | 27.13 ± 4.57 |
| SYDBAT Naming, score out of 30 | 26.29 ± 2.98 |
| Visual Reproduction II, score out of 43 | 29.34 ± 8.36 |
| Trail Making Test B, seconds | 65.48 ± 34.18 |
| Verbal Fluency, score | 46.02 ± 11.13 |
| Hooper Visual Organisation Test, score out of 30 | 23.98 ± 3.18 |
| **Neuroimaging** |  |
| BBB Water Exchange Rate, min<sup>-1</sup> | 117.73 ± 23.80 |
| Free Water, volume fraction | 0.07 ± 0.01 |
| Total ePVS Volume, mm<sup>3</sup> (median, IQR) | 3102.96 ± 1541.49 |
| Total ePVS, volume fraction | 0.006 ± 0.003 |
| BG ePVS, volume fraction | 0.009 ± 0.003 |
| WM ePVS, volume fraction | 0.006 ± 0.003 |
| Cerebral Blood Flow, mL/100g/min | 38.45 ± 8.41 |
| WMH Volume, mm<sup>3</sup> (median, IQR) | 529.41 ± 990.72 |
| WMH Volume, log +1 | 6.31 ± 1.56 |
| Cortical Thickness, cm | 2.44 ± 0.07 |
| Ventricle Volume, mm<sup>3</sup> | 22832.28 ± 13677.15 |
| Ventricle Volume, volume fraction | 0.014 ± 0.007 |
| Hippocampal Volume, mm<sup>3</sup> | 8034.67 ± 714.28 |
| Hippocampal Volume, volume fraction | 0.0052 ± 0.0005 |
| Free Water-corrected FA, degree of anisotropy | 0.48 ± 0.02 |
| **Plasma Biomarkers** |  |
| Aβ42/40 Ratio | 0.06 ± 0.01 |
| GFAP, pg/mL | 101.23 ± 43.61 |
| NfL, pg/mL | 18.7 ± 5.84 |
| pTau181, pg/mL | 21.22 ± 8.14 |
| pTau217, pg/mL | 0.37 ± 0.22 |

**Note:** Reported as mean ± standard deviation unless otherwise specified. Neuroimaging outcome measures reflect whole brain means within the grey matter, white matter, or both, as described further in the text.
Aβ = amyloid-beta; BBB = blood-brain barrier; BG = basal ganglia; CDR = clinical dementia rating; ePVS = enlarged perivascular space; FA = fractional anisotropy; GFAP = glial fibrillary acidic protein; HDL = high-density lipoprotein; LDL = low-density lipoprotein; MMSE = mini mental state exam; NfL = neurofilament light chain; pTau = phosphorylated tau; SYDBAT = Sydney Language Battery; WM = white matter; WMH = white matter hyperintensity.

We then examined relationships between construct composites usings Pearson’s partial correlations, adjusting for sex and intracranial volume. To assess associations of the construct composites scores with age and cognition, we performed linear regression analyses adjusting for sex, intracranial volume and additionally adjusted for age and education when assessing cognition.

## Results

Seventeen participants were excluded due to incomplete or poor-quality MRI scans, a further three were excluded due to incomplete biomarker analysis and a further two due to incomplete blood pathology results, leaving 127 participants available for the present study. The mean age was 67 years and 68% of the sample were women. A summary of cohort characteristics is in Table 1.

### Latent Constructs

The exploratory factor analysis revealed five latent constructs supported by the parallel analysis scree plots (eFigure 1) and measures of model fit (Tucker Lewis Index of Reliability = 0.991; RMSEA index = 0.011). The first factor, which we termed the “Brain & Vascular Health” construct, comprised greater hippocampal volume, greater BG ePVS, greater CBF, higher HDL cholesterol, lower ventricle volume and lower BMI. The second factor, termed the “Structural Integrity” construct, included greater cortical thickness, greater fwFA and greater BG ePVS. The third factor, the “Fluid Transport” construct, was defined by greater WM ePVS and higher Free Water. The fourth factor, the “AD Biomarkers” construct, comprised lower Aβ42/40 ratio, higher pTau181, and higher pTau217. The fifth factor, the “Neuronal Injury” construct, comprised higher GFAP and higher NfL (Table 2). BBB kw, WMH, LDL cholesterol and systolic BP did not meaningfully load onto any of the five factors (<0.30 loading across all).

**Table 2.**
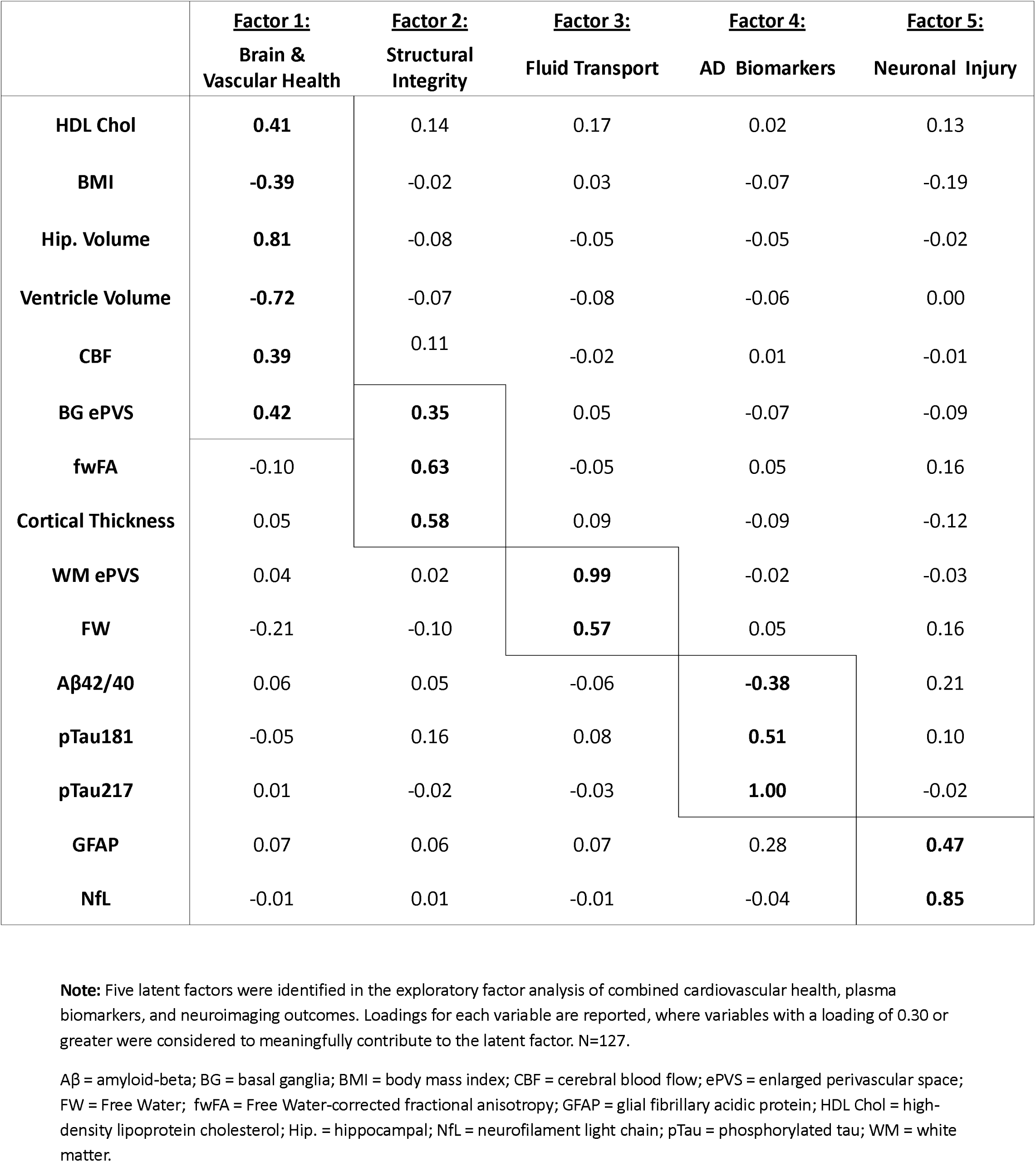
Latent Factors.

|  | **<u>Factor 1:</u>**<br>**Brain & Vascular Health** | **<u>Factor 2:</u>**<br>**Structural Integrity** | **<u>Factor 3:</u>**<br>**Fluid Transport** | **<u>Factor 4:</u>**<br>**AD Biomarkers** | **<u>Factor 5:</u>**<br>**Neuronal Injury** |
| --- | --- | --- | --- | --- | --- |
| **HDL Chol** | **0.41** | 0.14 | 0.17 | 0.02 | 0.13 |
| **BMI** | **-0.39** | -0.02 | 0.03 | -0.07 | -0.19 |
| **Hip. Volume** | **0.81** | -0.08 | -0.05 | -0.05 | -0.02 |
| **Ventricle Volume** | **-0.72** | -0.07 | -0.08 | -0.06 | 0.00 |
| **CBF** | **0.39** | 0.11 | -0.02 | 0.01 | -0.01 |
| **BG ePVS** | **0.42** | **0.35** | 0.05 | -0.07 | -0.09 |
| **fwFA** | -0.10 | **0.63** | -0.05 | 0.05 | 0.16 |
| **Cortical Thickness** | 0.05 | **0.58** | 0.09 | -0.09 | -0.12 |
| **WM ePVS** | 0.04 | 0.02 | **0.99** | -0.02 | -0.03 |
| **FW** | -0.21 | -0.10 | **0.57** | 0.05 | 0.16 |
| **Aβ42/40** | 0.06 | 0.05 | -0.06 | **-0.38** | 0.21 |
| **pTau181** | -0.05 | 0.16 | 0.08 | **0.51** | 0.10 |
| **pTau217** | 0.01 | -0.02 | -0.03 | **1.00** | -0.02 |
| **GFAP** | 0.07 | 0.06 | 0.07 | 0.28 | **0.47** |
| **NfL** | -0.01 | 0.01 | -0.01 | -0.04 | **0.85** |

**Note:** Five latent factors were identified in the exploratory factor analysis of combined cardiovascular health, plasma biomarkers, and neuroimaging outcomes. Loadings for each variable are reported, where variables with a loading of 0.30 or greater were considered to meaningfully contribute to the latent factor. N=127.
Aβ = amyloid-beta; BG = basal ganglia; BMI = body mass index; CBF = cerebral blood flow; ePVS = enlarged perivascular space; FW = Free Water; fwFA = Free Water-corrected fractional anisotropy; GFAP = glial fibrillary acidic protein; HDL Chol = high-density lipoprotein cholesterol; Hip. = hippocampal; NfL = neurofilament light chain; pTau = phosphorylated tau; WM = white matter.

Greater BG ePVS loaded onto the Brain & Vascular Health and Structural Integrity constucts, which went against our expectations. To better understand this finding, we conducted post-hoc analyses, described in the Supplement. Briefly, we conducted linear regressions investigating relationships between both BG region size and age with BG ePVS. We found that the pairing of BG ePVS with better health outcomes was not solely explained by basal ganglia region size and determined that BG ePVS is negatively associated with age (eFigure 2). Although these analyses do not rule out other potential methodological biases, it does suggest that our unexpected BG ePVS findings may have biological relevance. We consider the implication of these findings further in the Discussion, section “BG ePVS may be a contextual health marker”.

**Figure 2:**
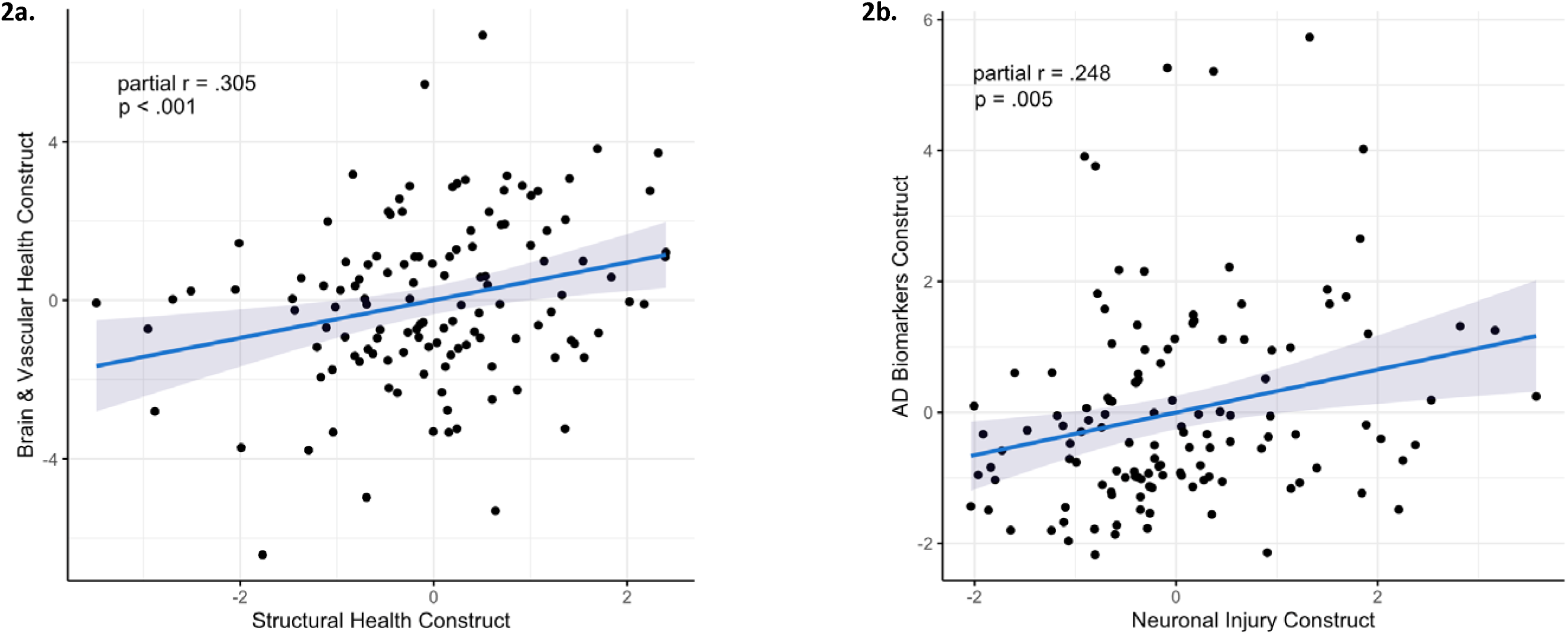
Correlations between Construct Composites. Pearson’s partial correlations adjusting for sex and intracranial volume revealed two statistically significant relationships between construct composites. a) a positive correlation between Brain & Vascular Health and Structural Health constructs. b) a positive correlation between Alzheimer’s disease (AD) Biomarkers and Neuronal Injury constructs.

### Correlations between Construct Composites

The Brain & Vascular Health and Structural Integrity constructs were positively correlated (*partial r*=.305, *t*=3.554, p<.001; Figure 2a), as were the AD Biomarker and Neuronal Injury constructs (*partial r*=.248, *t*=2.834, *p*=.005; Figure 2b). None of the other constructs were significantly correlated (eFigure 3). Since the correlation between the Brain & Vascular Health and Structural Injury constructs may be explained by both including the BG ePVS variable, we conducted a sensitivity analysis removing BG ePVS from the Brain & Vascular Health construct. We found that the correlation was attenuated but still statistically significant (*partial r*=.186, *t*=2.104, *p*=.037), suggesting that the shared inclusion of BG ePVS does not wholly explain this relationship.

**Figure 3:**
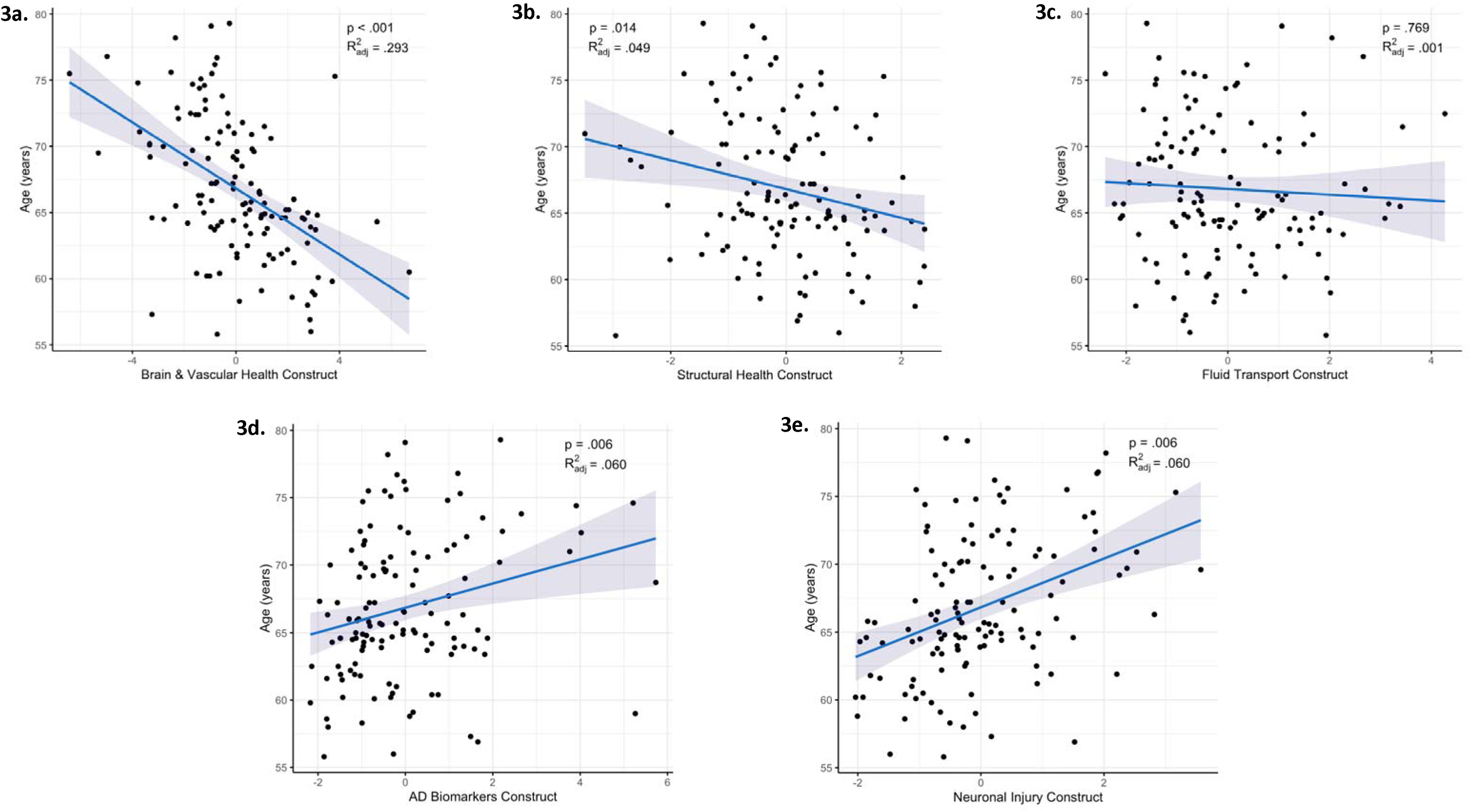
Associations between Construct Composites and Age. Multiple linear regression adjusting for sex and estimated intracranial volume. AD = Alzheimer’s disease.

**Figure 4:**
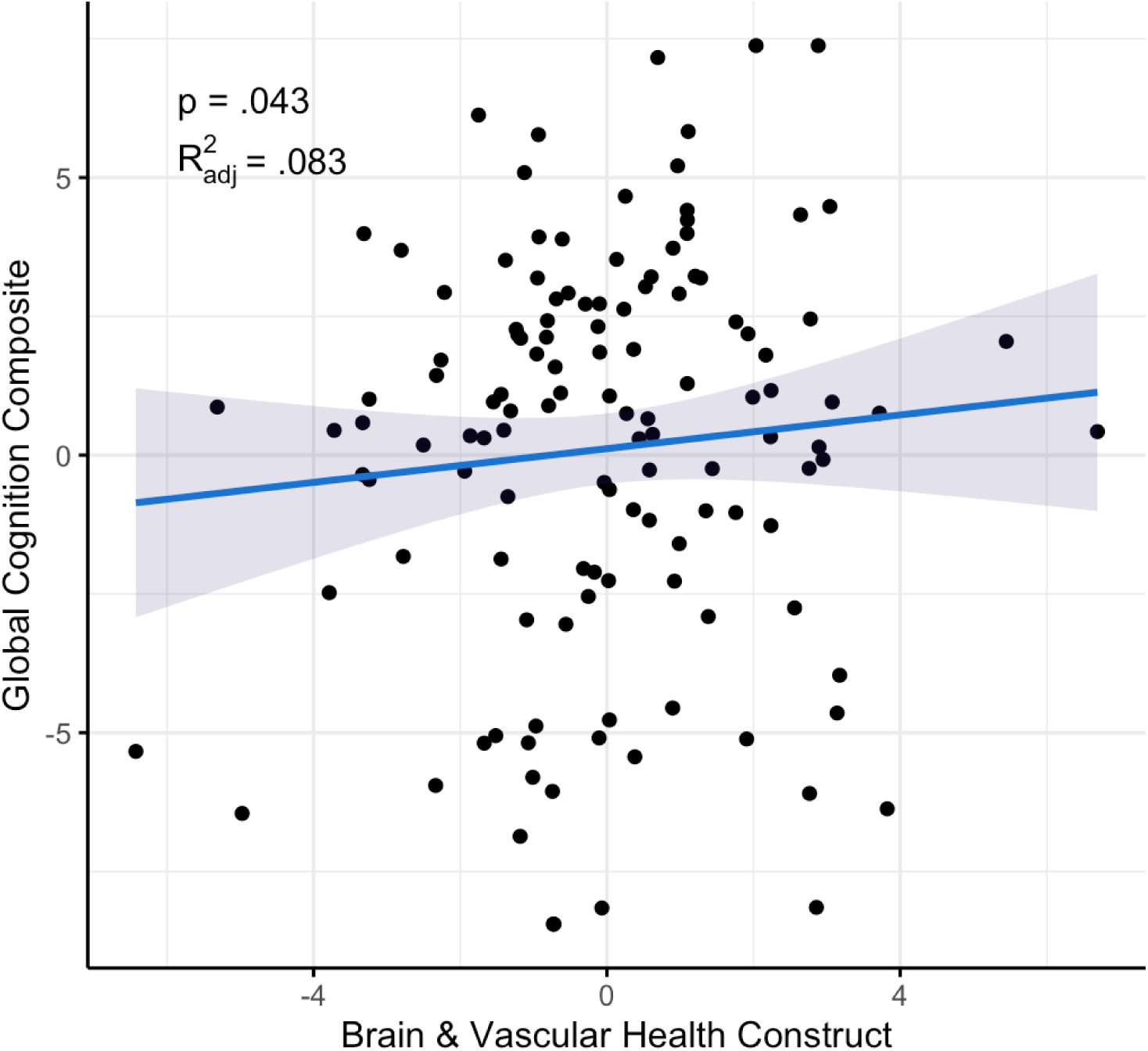
Positive Association between the Brain & Vascular Health Construct and Global Cognition. After adjustment for age, sex intracranial volume and years of education, a linear regression revealed a positive association between the Brain & Vascular Health construct and global cognition.

### Construct associations with Age

The Brain & Vascular Health (β=-0.70, SE=0.10, *p*<.001, R^2^ =.293) and Structural Integrity (β=-0.22, SE=0.09, *p*=.014, R^2^ =.049) constructs were negatively associated with age, while the AD Biomarkers (β=0.24, SE=0.09, *p*=.006, R^2^ =.060) and Neuronal Injury (β=0.39, SE=0.08, *p*<.001, R^2^ =.156) constructs were positively associated with age (Figure 3). The Fluid Transport construct was not significantly associated with age (β=-0.03, SE=0.10, *p*=.769, R^2^ =.001).

### Construct associations with Cognition

The Brain & Vascular Health construct was positively associated with global cognition (β=0.27, SE=0.13, *p*=.043, R^2^ =.083). However, the Structural Integrity (β=0.08, SE=0.09, *p*=.389, R^2^ =.057), Fluid Transport (β=-0.04, SE=0.10, *p*=.706, R^2^ =.052) and AD Biomarkers (β=-0.065, SE=0.09, *p*=.473, R^2^ =.055) and were not significantly associated with cognition. The Neuronal Injury construct had a nominal association with global cognition (β=-0.18, SE=0.09, *p*=.061, R^2^_adj_=.079). The associations between the constructs and individual cognitive tests are described in the supplement (eTable 3). Briefly, the Neuronal Injury construct was associated with Similarities (β=-0.22, SE=0.09, *p*=.019) and Visual Reproduction Delayed scores (β=-0.19, SE=0.09, *p*=.045), but none of the other constructs were associated with individual test scores.

## Discussion

We used multimodal data from MRI, biofluids and health assessments that are sensitive to neurovascular health and neurodegeneration to identify a set of underlying constructs related to aging and early brain pathology in a community cohort. We identified five latent constructs, which we labelled Brain & Vascular Health, Structural Integrity, Fluid Transport, AD Biomarkers and Neuronal Injury. Four of the five constructs (all except Fluid Transport) were associated with age, but only some constructs covaried with others (Brain & Vascular Health with Structural Integrity, and AD Biomarkers with Neuronal Injury). This suggests that while some of these constructs - and thus the biological systems they may represent - are interconnected, all of the constructs were distinct and largely separate. Furthermore, only one construct (Brain & Vascular Health) was significantly associated with global cognitive performance. These observations indicate that these constructs reflect relatively unique aspects of healthy and/or pathological brain aging and provide a set of biomarker composites that may have utility in disentangling different aspects of brain health. This work indicates that multimodal integration across traditional measures of brain health may offer a more nuanced approach to assessing the impacts of aging and pathology.

### Vascular Health and Structural Integrity

Our exploratory factor analysis identified a distinct “Brain & Vascular Health” construct that integrated markers of both neural and vascular integrity. This factor is characterised by variability in brain structure, such as relatively greater hippocampal volume and less ventricle enlargement (which serves as an indirect marker of reduced brain atrophy), in combination with indicators of more optimal vascular health, including greater CBF, greater BG ePVS and a reduced cardiovascular disease risk profile (lower BMI and higher HDL cholesterol levels). It was interesting that our agnostic approach identified a shared core construct across this broad set of measures together, as aspects of peripheral vascular and neurovascular health are often investigated separately in the context of aging despite their known dependencies^52^. This construct composite supports existing evidence that the health of the vasculature and brain are intricately linked^52^, and these measures may provide a more nuanced metric of health status when combined.

The observed associations between the Brain & Vascular Health construct, age and global cognition are particularly noteworthy. It is possible that our composite measure could serve as a proxy for overall brain health, conceptually akin to the metric of “brain age” derived from structural imaging. Moreover, its correlation with the separate “Structural Health” construct, comprising measures of white matter, cortical grey matter and deep grey matter structural integrity, further supports the idea that vascular health is a critical component in maintaining the broader architecture and function of the brain. Interestingly, the Brain & Vascular health construct was not significantly associated with factors representing “AD Biomarkers” or “Neuronal Injury”. This suggests that in a non-clinical community sample of older adults, the Brain & Vascular Health construct may primarily reflect vascular and structural changes associated with aging, which may be distinct from but potentially interact with AD pathology. For instance, even in the absence of neurodegenerative disease, cardiovascular risk factors tend to increase while brain volumes and CBF tend to decrease with age^53–55^. Extrapolating further, the identification of this construct supports growing evidence that maintaining optimal vascular health may offer a degree of protection against age-related brain atrophy, independent of the mechanisms driving AD pathology. Although optimal cardiovascular health is known to protect against dementia^17^, and both AD and vascular brain injury are frequently comorbid, the extent to which vascular health mitigates the risk of AD pathology remains unclear. The observed patterns suggest that age-related vascular decline may follow a distinct trajectory from AD, a hypothesis that warrants further exploration in longitudinal studies. Research and intervention trials investigating whether maintaining optimal vascular health can mitigate the earliest seedings of AD would significantly advance our understanding of vascular effects on neurodegenerative pathology.

### Fluid Stagnation

We identified a fluid stagnation construct characterized by greater extracellular Free Water and greater WM ePVS volume. This construct is consistent with previous research linking these MRI markers to fluid transport dysfunction, particularly in the extracellular fluid of the brain parenchyma^8,56,57^. Interestingly, although Free Water is thought to be a broad marker of both fluid transport stagnation and microstructural integrity, fw-FA did not load on this factor. This supports the notion that, in the context of increased WM ePVS, Free Water represents fluid-related dysfunction. Notably, we did not find a correlation between this construct with either the AD Biomarkers or Neuronal Injury constructs that comprised plasma biomarker measures, highlighting the potential of MRI markers to capture aspects of dysfunction or breakdown that these biomarkers may not fully reflect.

The Fluid Transport construct was the only construct not associated with age in our analysis. In our previous work, we found that although Free Water was associated with age, the inter-relationship between Free Water and ePVS was not modified by age^7^. This finding has been replicated in a more recent study which showed that the ratio between WM ePVS and Free Water remains stable through age^58^. We postulated that this consistent relationship between WM ePVS and Free Water represents the interconnected compartments of the fluid transport system (within the neurovascular unit and extracellular parenchyma), which does not decouple through aging. Given this, the Fluid Transport construct may provide an avenue to assess fluid transport dysfunction independent of age-related changes. Future research is needed to investigate whether this Fluid Transport construct may be a robust marker for changes in the brains waste clearance system.

### AD Biomarkers and Neuronal Injury

Our findings revealed distinct constructs for AD biomarkers and neuronal integrity. The AD Biomarkers construct reflected the emergence of hallmark amyloid and tau pathology, while Neuronal Injury construct encompassed biomarkers of astrocyte reactivity or neuronal breakdown (GFAP) and axonal degeneration (NfL). Interestingly, these constructs were not correlated with one another in our data and may represent distinct biological processes. This suggests that while cellular breakdown may accompany AD pathology, it could represent a secondary process to amyloid and tau accumulation. Research has demonstrated that GFAP and NfL changes are not exclusive to AD, as their elevations are observed across various states of brain health, including acquired injury^59,60^ and other neurodegenerative or neuroinflammatory conditions^61,62^.

Longitudinal studies of AD biomarker levels have shown that plasma pTau217 seems to be particularly sensitive to the earliest processes related to AD^63,64^ and is also more predictive of abnormal amyloid levels on PET imaging than either plasma pTau181 or Aβ42/40^65,66^. Thus, it is perhaps unsurprising that pTau217 had the highest loading on the identified AD Biomarkers construct. Prior research has proposed that plasma pTau217 could be related to both amyloid accumulation and tau phosphorylation processes in AD^65,67^, which is also reinforced by our construct composite including measures of pTau217, pTau181 and Aβ42/40 together.

Taken together, the weighted combination of plasma pTau217, pTau181 and Aβ42/40 seemed to describe elevated AD Biomarkers even in our community-based cohort, which is in accordance with previous studies^68^. Future research is encouraged to explore whether weighted composites might better detect the early emergence of AD neuropathology as compared to individual markers alone.

### BG ePVS may be a contextual health marker

In this study, greater BG ePVS volume fraction loaded positively onto the Brain & Vascular Health and Structural Integrity factors alongside measures such as greater hippocampal volume, greater cerebral blood flow and higher white matter microstructural integrity. Our further analyses indicated that is not merely an effect of available brain area and is likely a true biological finding within our sample. Notably, WM ePVS but not BG ePVS paired with increased Free Water in the Fluid Stagnation construct (in fact, Free Water loaded slightly negatively on factors that involving greater BG ePVS), emphasising regional differences in ePVS interpretation in our data.

It is important to consider that the BG ePVS quantified in this analysis may differ from those described in visual scoring approach, such as typical for diagnosis of cerebral small vessel disease^69,70^. Quantified ePVS volume and qualitatively rated ePVS may represent slightly different measures of perivascular space enlargement, where rating of severity emphasises frequency and quantification of volume emphasises dilation or augmentation. Moreover, the automatic segmentation method we employed has been demonstrated to have improved sensitivity and accuracy for BG ePVS compared to other available algorithms^32^ for quantifying ePVS volume. Researchers are encouraged to consider these factors when interpreting seemingly discrepant findings across research studies.

The interpretation of greater BG ePVS volume as a marker of good health is novel and challenges existing interpretations of MRI-visible perivascular spaces as purely pathological imaging features^70,71^. Recent literature reviews and meta-analyses have highlighted that overall, the associations between BG ePVS, cognition^72^ and AD^34^ are ambiguous, partly due to heterogeneity of methodology (whether ePVS were quantified as count or volume), cohort (comorbidities or disease diagnosis) and adjustment for confounders. In light of these inconsistent findings, researchers have postulated that the enlargement of perivascular spaces is part of compensatory or neuroprotective mechanisms amidst cognitive decline and pathology^72–74^.

It is possible that in the context of healthy aging, an elevated volume of BG ePVS may reflect more efficient fluid transport, perhaps through normal adaptive mechanisms that respond to metabolic activity – particularly since the BG structures are highly metabolically active^75^. This may be especially true for adults aged 50 to 80, as previous cross-sectional studies have shown a relative plateau in ePVS increase during this life period^76,77^, potentially allowing better detection of variability due to factors other than age. This aligns with the notion that perivascular spaces themselves are essential for fluid dynamics and waste clearance rather than inherently pathological^78^. While findings in the context of healthy aging are limited, previous research has reported similar effect directions: one study showed that increased BG ePVS was associated with decreased tau deposition in the brain in cognitively impaired participants and in those with genetic AD risk (apolipoprotein E4 carriers)^79^, whereas another demonstrated that older adults with better sleep quality had larger BG ePVS^80^. We postulate that in those without overt neurological or vascular disease (such as the co-existence of other imaging features of cerebral small vessel disease), greater BG ePVS volume may reflect increased flow or clearance within the perivascular space within the deep gray matter, and that the interpretation of BG ePVS may depend on the broader context of brain state and associated measures. Future research should investigate whether greater BG ePVS volume might represent more optimal vascular health in aging under specific conditions, while remaining cautious about the potential duality of ePVS in health and disease.

### Study Strengths & Limitations

This study was the first, to our knowledge, to combine multimodal imaging and fluid biomarker data to derive latent constructs that provide insight into underlying mechanisms of brain health. We analysed a broad scope of high-quality neuroimaging measures, plasma biomarkers and cardiovascular risk factors, implemented advanced neuroimaging analysis techniques and utilised gold-standard assays for fluid biomarker analysis. By studying a non-clinical community cohort, we were able to establish a framework for investigating separable components of brain health in an older population. This offers a reference point for comparison with neurological conditions, where overlapping pathologies and more pronounced dysfunction may obscure distinctions between vascular, structural and neurodegenerative processes.

Despite these strengths, there are limitations to consider. We did not find associations between many of our identified constructs and cognition, despite previous literature linking some individual components, such as cortical thickness, Aβ42/40, pTau181, pTau217, GFAP and NfL, with poorer cognition through a broad cognitive spectrum^81,82^. Given that we exclude participants with moderate-severe cognitive impairment from the BACH study, it is possible that different relationships with cognition would emerge in the setting of clinical cognitive impairment. Similarly, while our findings provide a reference framework for healthy aging and potentially pre-clinical and prodromal Alzheimer’s disease, the generalisability to other clinical or non-clinical populations or those with advanced pathology remains uncertain. Future research could address these gaps by integrating longitudinal designs, validating constructs against histopathological or other gold-standard measures, and expanding the scope to include diverse cohorts with varying degrees of brain pathology and cognitive impairment.

## Conclusion

This study advances our understanding of brain health and disease by identifying distinct latent constructs through the integration of multimodal imaging and fluid biomarkers. The Brain & Vascular Health, Structural Integrity, Fluid Transport, AD Biomarkers, and Neuronal Injury constructs provide a comprehensive framework for examining the complex interplay between neurodegeneration, fluid transport, and vascular health in aging. Our findings highlight that while these constructs collectively contribute to our understanding of brain health, they reflect distinct biological processes with unique mechanisms. By establishing a reference for healthy aging, we demonstrated the utility of using multiple biomarkers to probe systems of neurovascular and brain health that are not well captured by a single measure. Furthermore, our approach provides a framework that could be adopted by future studies to explore brain changes through aging and neurodegenerative disease continuums. Our findings reinforce the importance of considering vascular and fluid transport dysfunction alongside traditional AD biomarkers to refine our understanding of aging and disease progression. Future research should validate these constructs in diverse populations, assess their longitudinal trajectories in aging, and evaluate their predictive value for cognitive decline and dementia risk.

## Acknowledgments

We would like to thank the Brain and Cognitive Health (BACH) cohort study participants and their families for their invaluable time and contributions to this research.

## Statements and Declarations

### Author Contributions

ER, ML, MPP and IHH were involved in study conception. ER, KF, TC, SY, MC, LV, TJO, ML, IHH and MPP were involved in cohort project design and data collection. ER conducted processing and analysis of MRI measures in consultation with BS, XS, DJJW, ML and IHH. MAS, WTO and SJM conducted analysis of biofluid markers. ER conducted statistical analysis. ER, IHH and MPP prepared the manuscript draft. All authors read and approved the final manuscript.

