## Supplementary Material for "Factor Analysis of Multimodal MRI, Biofluid and Vascular Health Biomarkers Reveals Latent Constructs of Brain Health"

**Correspondence:**

Assoc/Prof. Matthew P. Pase

**Principal Components Analysis of Cognitive Tests**

We conducted a principal components analysis to determine tests that best represented global cognitive function to form our primary cognition outcome variable. Firstly, we used parallel analysis scree plots to guide the number of factors that best suit the data, where a single-factor, two-factor, three-factor or four-factor models were deemed appropriate. Across all factor solutions, the first principal component (PC1) consistently emerged as a robust measure of global cognition, comprising seven of the nine cognitive outcomes. The four-factor model (**eTable 1**) produced the most optimal fit metrics, had high interpretability and was therefore retained for further analyses (model comparison in **eTable 2**).

To evaluate the internal consistency of the cognitive tests contributing to the global cognition factor (PC1), we calculated Cronbach's alpha using the factor loadings form the four-factor PCA model. All cognitive tests that loaded strongly onto PC1 (>0.45) were included in the analysis. Cronbach's alpha for PC1 was 0.77 (Feldt's 95% CI = [0.70, 0.83]), and no single item would improve alpha if removed, suggesting internal consistency across all included tests.

**eTable 1. Principal Components Analysis of Cognitive Tests: Four-factor model loadings**

| | PC1 | PC2 | PC3 | PC4 |
|---|---|---|---|---|
| **Prose Passages Delayed** | **0.50** | **0.49** | -0.32 | 0.32 |
| **Similarities** | **0.67** | 0.41 | -0.12 | 0.10 |
| **SYDBAT Naming** | **0.76** | 0.01 | **0.42** | -0.01 |
| **Visual Repro. Delayed** | **0.64** | 0.30 | 0.08 | 0.02 |
| **TMT A*** | -0.44 | **0.51** | **0.43** | 0.43 |
| **TMT B*** | **-0.72** | 0.15 | 0.33 | 0.06 |
| **Verbal Fluency** | **0.63** | 0.03 | 0.09 | -0.44 |
| **HVOT** | **0.53** | -0.41 | **0.59** | 0.15 |
| **TASIT** | 0.32 | **-0.61** | -0.24 | **-0.58** |

**Note:** *Lower values indicate better performance.
Loadings >0.45 were considered to strongly load onto a given factor.
SYDBAT = Sydney Language Battery ; Visual Repro. = Visual Reproduction; TMT = Trail Making Test; HVOT = Hooper Visual Organisation Test; TASIT = The Awareness of Social Inference Test.

**eTable 2. Principal Components Analysis of Cognitive Tests: Model Comparison**

| Model | RMSR | Chi-Square ($\chi^2$) | $\chi^2$ p-value | Variance Explained* | Mean Item Complexity |
|---|---|---|---|---|---|
| One-factor | 0.12 | 127.60 | $5.4e^{15}$ | 83% | 1.0 |
| Two-factor | 0.11 | 117.64 | $3.1e^{16}$ | 85% | 1.5 |
| Three-factor | 0.10 | 91.29 | $2.8e^{14}$ | 88% | 1.9 |
| Four-factor | 0.09 | 71.23 | $2.3e^{13}$ | 91% | 2.4 |

**Note:** RMSR = root mean square of residuals.
*Percentage of variance explained reflects the off-diagonal variance in the correlation matrix.
N=127

**1a.**

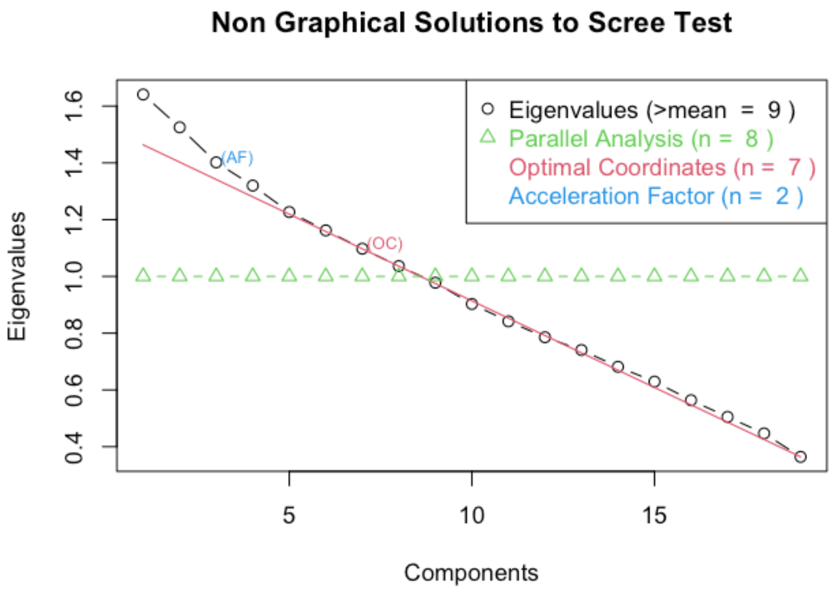


**1b.**

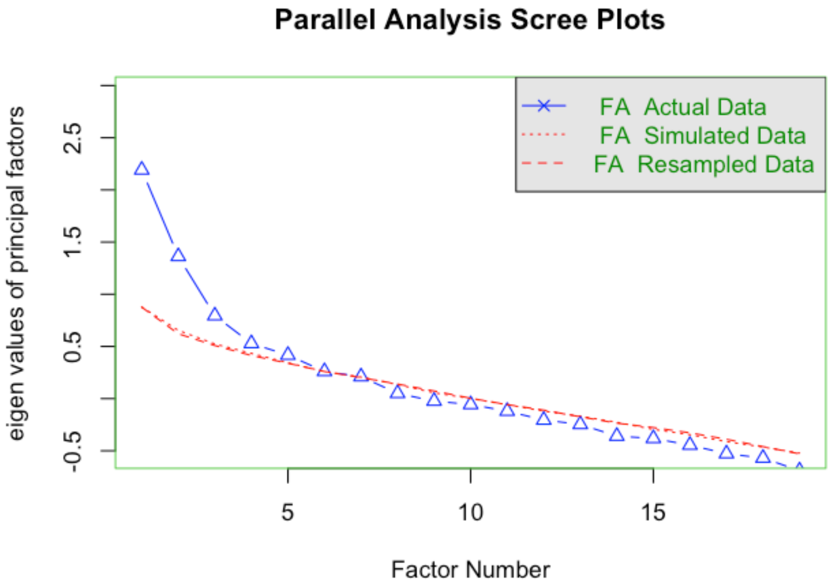


**eFigure 1: Exploratory Factor Analysis (EFA) Scree Plots**

Scree plots were used to determine the numbers of factors to retain. While the traditional 'elbow' method was less clear, five factors were retained as they met criterion below and maintained conceptual coherence.

**a. Non-Graphical Solutions to the Scree Test:** This plot compares multiple factor retention methods. Parallel analysis (green) suggested 8 factors, optimal coordinates (red) suggested 7, and the acceleration factor method (blue) suggested 2. A five-factor solution was supported as a balance between these methods and theoretical interpretability.

**b. Parallel Analysis Scree Plot:** This plot offers a robust refinement of the scree test findings. The retainment of n-factors was supported if their eigenvalues (blue) exceeded those of the simulated (dotted red) and resampled (dashed red) data. This method also supported a five-factor solution.

**Basal Ganglia ePVS Post-hoc Analysis**

We observed that greater basal ganglia (BG) enlarged perivascular space volume (ePVS) loaded onto the Brain & Vascular Health and Structural Health factors, which went against our expectations. To better understand this finding, we conducted post-hoc analyses, first exploring potential methodological bias. While we analysed ePVS as a volume fraction to minimise the effect of individual region volume variability, it is possible that those with larger brain volumes (or less atrophy) have a larger available area for ePVS to occur. Linear regressions adjusting for age, sex and intracranial volume confirmed a significant positive relationship between BG region volume and total BG ePVS volume (β=0.72, SE=0.11, $p$<.001, $R^2_{adj}$=.358), with a similar but attenuated effect for ePVS volume fraction (β=0.54, SE=0.12, $p$<.001, $R^2_{adj}$=.301). Interestingly, while previous cross-sectional studies report that ePVS volumes increase with age[1,2], we found that BG ePVS volume fraction *decreased* with age across our cross-sectional cohort (β=-0.39, SE=0.08, $p$<.001, $R^2_{adj}$=.151). There was a significant interaction effect between ePVS and BG region volume, where BG ePVS volume fraction instead increased with age in those with the very largest BG region volumes within our data (interaction p=.010), but decreased with age in the remainder of the cohort (**eFigure 2**). This demonstrates that the pairing of BG ePVS with better health outcomes and the negative relationship between BG ePVS and age is not solely explained by region size effects. Although this does not completely rule out other potential methodological biases, it does suggest that our BG ePVS findings may have biological relevance.

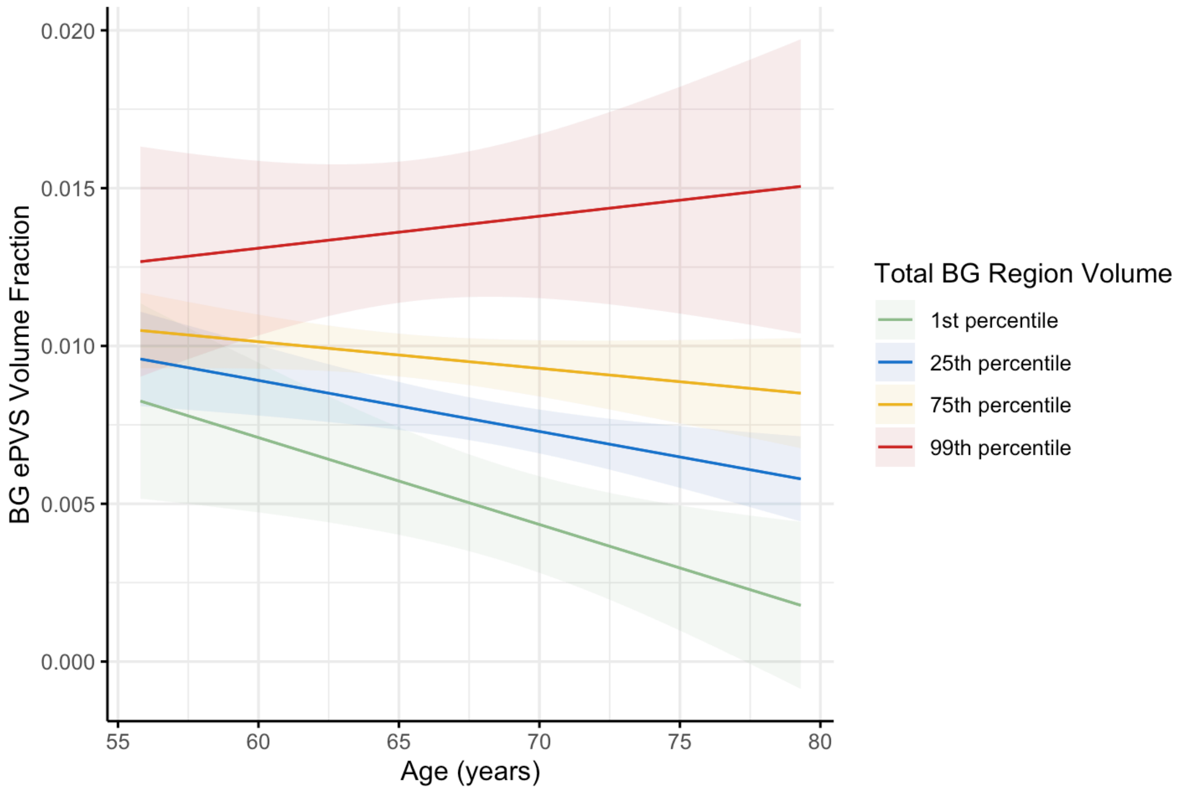


**eFigure 2: Interaction effect between BG region volume, BG ePVS volume fraction and age.**

A linear regression model adjusting for sex and intracranial volume tested the interaction effect of total basal ganglia (BG) region volume on the association between BG enlarged perivascular space (ePVS) volume fraction and age.

Although there was a significant interaction effect, BG ePVS volume fraction increased with age in only those with the very largest BG region volumes within our data (at approximately the 99$^{th}$ percentile) and decreased with age for the remainder of the cohort.

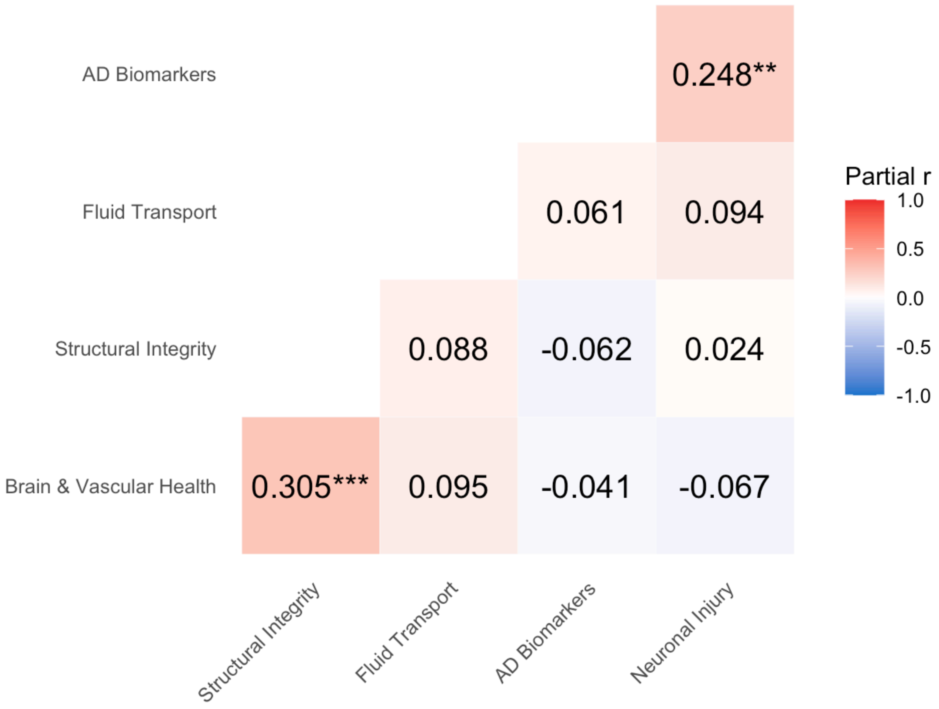


**eFigure 3. Correlations between Construct Composites**

Pearson's partial correlations between each of the construct composites, adjusting for sex and intracranial volume. AD = Alzheimer's disease.
***p*<.01, ****p*<.001.

**eTable 3. Relationships between Construct Composites and Individual Cognitive Tests**

| | Factor 1: Brain & Vascular Health | | Factor 2: Structural Integrity | | Factor 3: Fluid Transport | | Factor 4: AD Biomarkers | | Factor 5: Neuronal Injury | |
|---|---|---|---|---|---|---|---|---|---|---|
| | β (SE) | *p* | β (SE) | *p* | β (SE) | *p* | β (SE) | *p* | β (SE) | *p* |
| **Prose Passages Delayed** | 0.150 (0.140) | .287 | 0.091 (0.093) | .327 | -0.015 (0.098) | .878 | -0.059 (0.093) | .527 | -0.185 (0.097) | .060 |
| **Similarities** | 0.240 (0.135) | .073 | -0.037 (0.089) | .676 | -0.089 (0.094) | .345 | -0.119 (0.089) | .182 | **-0.219 (0.092)** | **.019** |
| **SYDBAT Naming** | 0.141 (0.139) | .312 | 0.016 (0.092) | .865 | -0.066 (0.097) | .498 | 0.024 (0.093) | .798 | -0.035 (0.098) | .722 |
| **Visual Reproduction Delayed** | 0.074 (0.134) | .583 | 0.093 (0.088) | .296 | 0.047 (0.094) | .617 | -0.040 (0.089) | .654 | **-0.188 (0.093)** | **.045** |
| **Trail Making Test B*** | -0.212 (0.132) | .111 | 0.002 (0.088) | .983 | -0.025 (0.093) | .790 | -0.084 (0.088) | .343 | 0.159 (0.092) | .087 |
| **Verbal Fluency** | 0.255 (0.135) | .061 | 0.046 (0.090) | .614 | -0.083 (0.095) | .383 | -0.036 (0.091) | .692 | -0.054 (0.096) | .570 |
| **Hooper Visual Organisation Test** | -0.042 (0.140) | .766 | 0.074 (0.092) | .425 | 0.042 (0.098) | .664 | 0.141 (0.092) | .129 | -0.033 (0.098) | .737 |

**Note:** *Lower values indicate better performance.

Multiple linear regressions between each cognitive outcome and factor constructs. All models adjusted for age, sex, estimated intracranial volume and education. AD = Alzheimer's disease; SYDBAT = Sydney Language Battery.
